# Endogenous neural stem cells modulate microglia and protect from demyelination

**DOI:** 10.1101/2020.06.18.158782

**Authors:** Béatrice Brousse, Karine Magalon, Fabrice Daian, Pascale Durbec, Myriam Cayre

## Abstract

In response to corpus callosum (CC) demyelination, subventricular zone-derived neural progenitors (SVZdNP) are mobilized and generate new myelinating oligodendrocytes. Here, we examine the putative immunomodulatory properties of endogenous SVZdNP during demyelination in the cuprizone model. We observed that SVZdNP density is higher in the lateral and rostral CC regions that show weaker demyelination and is inversely correlated with activated microglia density and pro-inflammatory cytokines levels. Single-cell RNA-sequencing further revealed CC areas with high SVZdNP mobilization are enriched in a microglial cell subpopulation with immunomodulatory signature. We identified ligand/receptor couple MFGE8 (milk fat globule-epidermal growth factor-8)/integrin β3 as a ligand/receptor couple implicated in SVZdNP/microglia dialog. MFGE8 is highly enriched in immature SVZdNP mobilized to the demyelinated CC and promotes myelin debris phagocytosis in vitro. Altogether these results demonstrate that beyond their cell replacement capacity endogenous progenitors display immunomodulatory properties highlighting a new role for endogenous SVZdNP in myelin regeneration.

## INTRODUCTION

Myelin regeneration has been observed in the brain of patients with multiple sclerosis (MS) but is not always effective and exhibits a very high variability among patients (Patrikios et al. 2006, Albert et al. 2007). In rodents, spontaneous remyelination after experimentally-induced demyelination is a very efficient process that can involve both parenchymal oligodendrocyte progenitor cells (pOPC) (Franklin et al. 1997, Gensert and Goldman 1997, Zawadzka et al. 2010) and adult neural stem/progenitor cells derived from the subventricular zone (SVZdNP) (Nait-Oumesmar et al. 1999, Picard-Riera et al. 2002, Menn et al. 2006, Aguirre et al. 2007) can participate to this repair process. In physiological conditions, neural stem cells (NSC) in the adult SVZ divide slowly and asymmetrically to give rise to actively proliferating progenitors (called “C cells) which in turn mainly produce neuroblasts and, at much lower rate, OPC (Doetsch et al. 1999, Menn et al., 2006). Demyelination induction in the periventricular white matter leads to a 4 fold increase in OPC production in the SVZ (Menn et al. 2006). Although the contribution of SVZdNP to spontaneous remyelination has long been considered negligible compared to pOPC, two independent studies uncovered their unexpectedly high and regionalized mobilization during cuprizone-induced demyelination in mice (Xing et al. 2014, Brousse et al. 2015).

Several studies using NSC grafts in injured rodent brain revealed a bystander effect of NSC independent of cell replacement, via the production of neurotrophic factors (Goldberg et al. 2015, Zuo et al. 2015), and immunomodulatory properties. NSC can dialog with blood-born infiltrating T cells and with microglial cells (Pluchino et al. 2005, Pluchino et al. 2009, Martino et al. 2011, Martino et al. 2011, Cusimano et al. 2012, Zhang et al. 2016).

Microglia plays key roles in demyelination/remyelination events (Chu et al., 2018). Activated microglia can mediate cell damage via cytokines and nitric oxide production. However, it is also involved in phagocytosis and removal of myelin debris which is a prerequisite of successful remyelination (Kotter et al. 2006, Lampron et al. 2015, Poliani et al. 2015). Consecutive to demyelination insults, microglial cells are activated and can adopt different phenotypes from M1 (pro-inflammatory) to M2 (immuno-modulatory) with many possible intermediate states (Miron et al. 2013, Lloyd et al. 2019, Vogel et al. 2013, Peferoen et al. 2015). Several studies suggest that NSC can promote the polarization of microglial cells toward the M2 phenotype. However these properties were either demonstrated after transplantation of hundreds of thousands (or even millions) of NSC (Gao et al. 2016, Marteyn et al. 2016), or *in vitro* in co-culture experiments (Liu et al. 2013, Wu et al. 2014).

Here, we examined whether and how endogenous SVZdNP spontaneously recruited to demyelinated lesions can also act through immunomodulation. We used the cuprizone model which lacks immune blood cell infiltration to focus on the interaction between endogenous SVZdNP and microglial cells. Analysis of CC areas with high or low SVZdNP recruitment allowed us to correlate the presence of SVZdNP with demyelination level and inflammatory signature. Single-cell RNA-sequencing of microglial cells sorted from these different CC areas, revealed clusters of cells enriched in M1 or M2 genes, suggesting an immunomodulatory phenotype in a population of microglial cells located in areas where SVZdNP are abundant. Finally we identified MFGE8 as a candidate factor secreted by SVZdNP for improving microglia phagocytotic properties.

## RESULTS

### Cuprizone-induced demyelination and SVZdNP mobilization are regionalized within the corpus callosum

Cuprizone is known to trigger diffuse demyelination throughout the brain. As expected, 3 weeks after the onset of cuprizone treatment, CC myelin content (assessed by GFP signal in PLP-GFP mice) decreased, reaching a minimum after 5 weeks of treatment, and then progressively recovered after return to normal diet (Fig1A, B). Demyelination extent along the rostro-caudal and medio-lateral axes of the CC was examined (Fig1C-E). In control mice not exposed to cuprizone, myelin content in CC is slightly higher medially (GFP ratio medial/lateral= 1.13±0.12) and rostrally (GFP ratio caudal/rostral= 0.85±0.04) (Fig1E). If demyelination proceeded homogeneously in the CC these ratios should remain constant during cuprizone exposure. But we observe a different scenario: the drop of medial/lateral fluorescence ratio from 1.13±0.12 to 0.82±0.06 (p=0.02) indicates that demyelination is clearly more pronounced in medial areas as soon as 3 weeks of cuprizone exposure (W3). After 5 weeks of cuprizone feeding (W5), demyelination becomes also heterogeneous along the rostro-caudal axis, caudal areas being more affected (32% reduction of caudal/rostral GFP ratio, from 0.85±0.04 to 0.58±0.05; p=0.03). These results show that both rostral and lateral CC are less affected by cuprizone treatment. After cuprizone removal and remyelination, regional differences in myelin content return to control levels (Fig1E).

**Figure 1:**
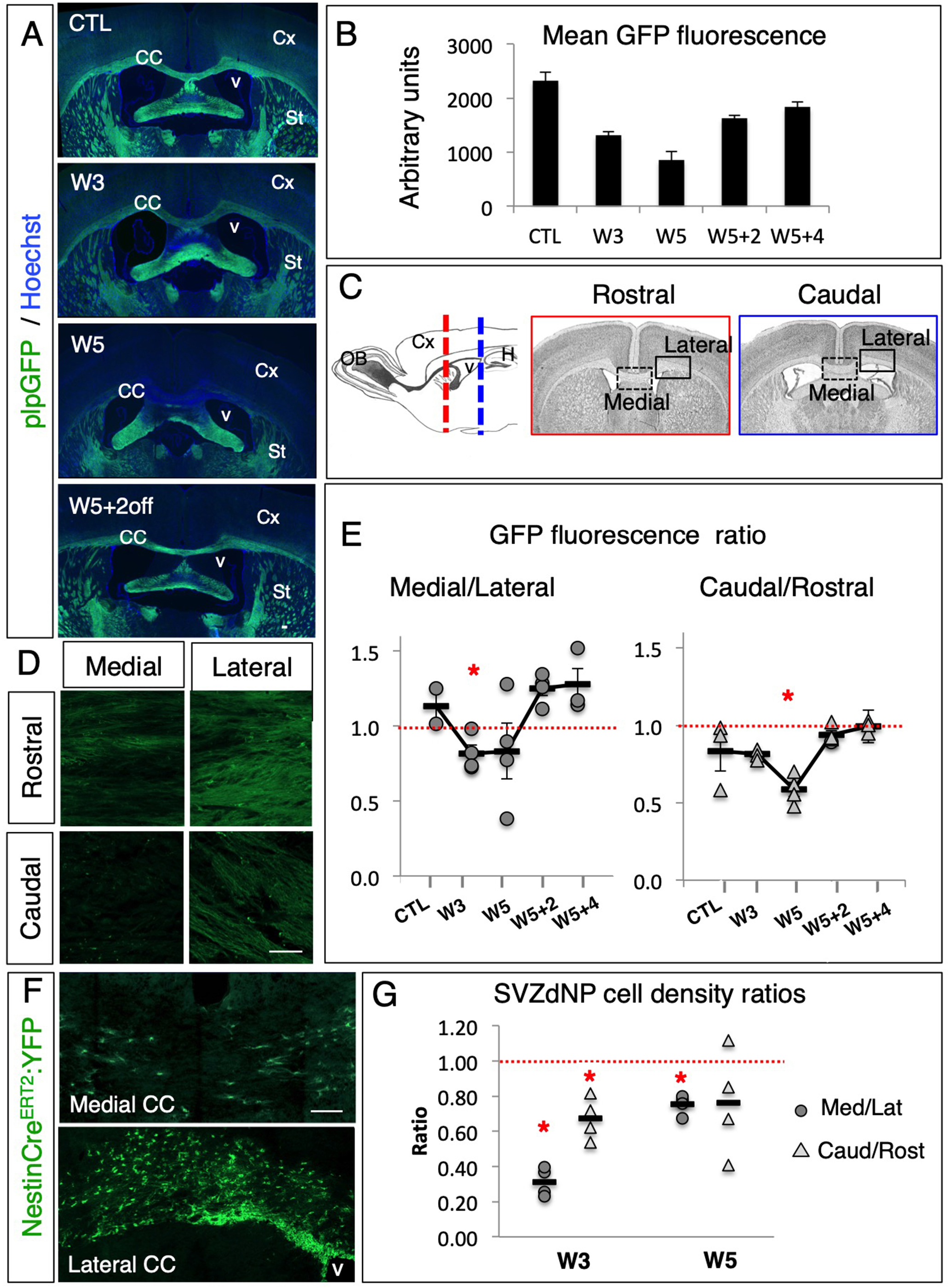
Demyelination and SVZdNP mobilization are regionalized within the corpus callosum following cuprizone intoxication. A,B: Illustration (A) and quantification (B) of demyelination and remyelination in plpGFP mice fed with cuprizone (mean ± sem; n=4 mice per time point). C: Location of CC areas analyzed. D: GFP signal in plpGFP mice in the different CC areas after 5 weeks cuprizone feeding. E: Quantification of GFP signal in medial relative to lateral CC (left panel) and in caudal relative to rostral CC (right panel) within each mouse during cuprizone-induced demyelination and remyelination. Ratio=1 means equal myelin content in medial and lateral CC (left panel) or in rostral and caudal CC (right panel). Black bars show the means, n=3 to 4 mice per time point. F,G: Illustration (F) and quantification (G) of SVZdNP mobilization in medial and lateral CC using NestinCre^ERT2^:YFP mice. SVZdNP recruitment is higher in lateral than medial CC, and in rostral than in caudal CC, as indicated by the mean ratio <1. Black bars show the means, n=4 mice per time point. Scale bars: 100 μm.

Having previously shown that SVZdNP preferentially contribute to the remyelination in the rostral CC (Brousse et al. 2015); we traced SVZdNP using NestinCre^ERT2^:rosaYFP transgenic mice and examined their distribution along both rostro-caudal and medio-lateral axes and calculated density ratios in each animal. In control mice, SVZdNP are virtually absent in CC (Brousse et al., 2015). As soon as W3, SVZdNP are preferentially recruited in the rostral and lateral CC. The differential recruitment is more pronounced along the medio-lateral axis with very few cells present in the medial CC (ratio medial/lateral = 0.31±0.04) (Fig1F,G). Such regionalized distribution of SVZdNP is maintained, although to a lesser extent, at W5 (0.75±0.03 and 0.76+0.15 for medial/lateral and caudal/rostral ratios respectively) (Fig1G).

These results show that the SVZdNP are recruited to the demyelinated CC with a regionalized pattern. They are preferentially recruited in the rostro-lateral CC just above the SVZ niche. Interestingly, these areas correspond to those less affected by cuprizone-induced demyelination (Fig1D, E), suggesting that these areas may be either protected from demyelination and/or start to be remyelinated faster even in presence of cuprizone.

Thus, we checked if the early differentiation of SVZdNP in mature oligodendrocytes (OLG) could explain the lower loss in myelin content in rostral and lateral CC during demyelination. At W3, although 9.8 ± 1.2 % of SVZdNP already differentiated into mature (CC1+) OLG, the cells do not yet adopt the morphology of myelinating OLG (Fig2A); at the end of cuprizone treatment (W5) 30.2 ± 3.0 % SVZdNP expressed CC1 and these cells began to exhibit typical myelin segments (Fig2B). This morphology is even more obvious after cuprizone removal (W5+2; Fig2C).

**Figure 2:**
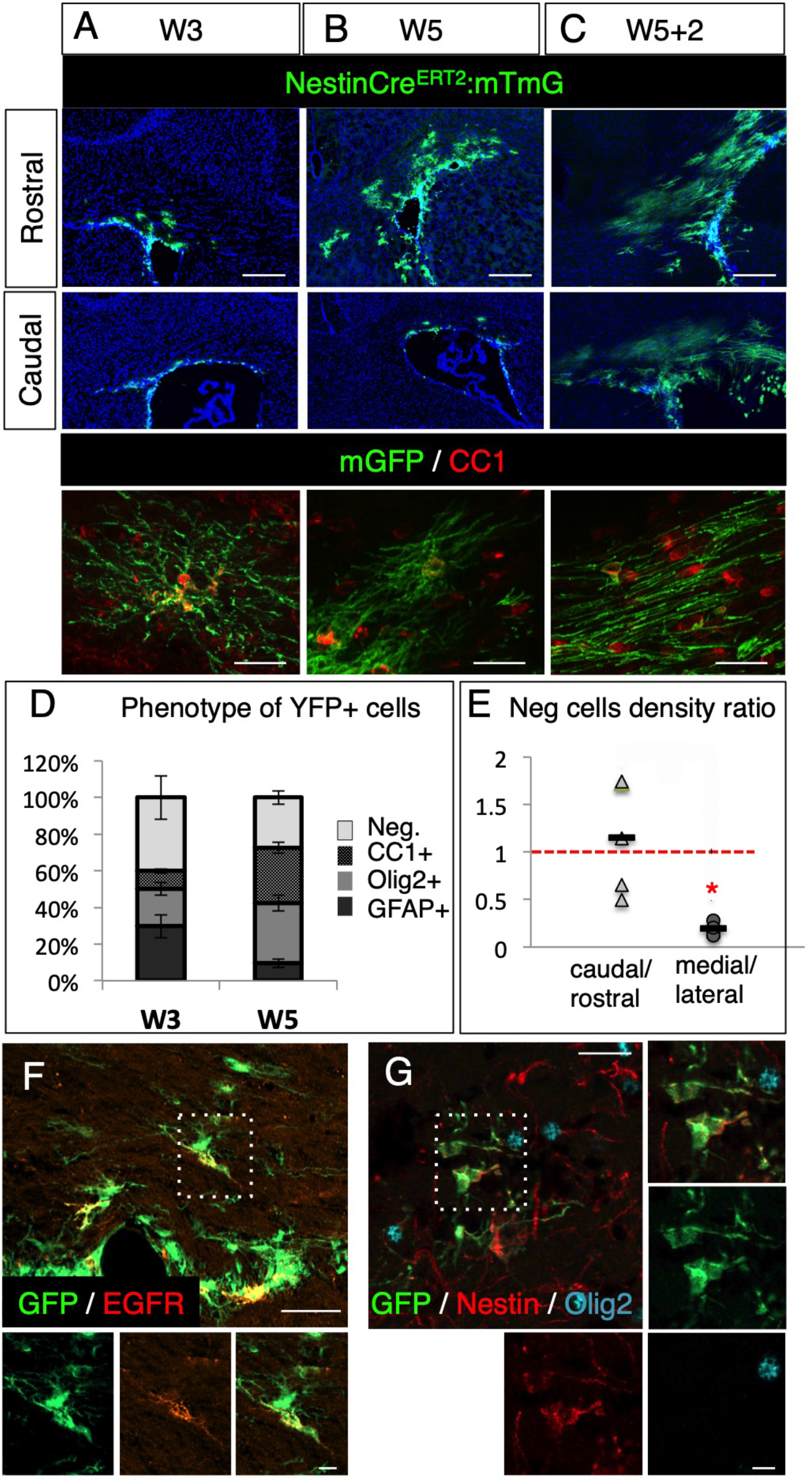
A significant proportion of SVZdNP remain undifferentiated in the demyelinating corpus callosum. A-C: Illustrations of remyelination by SVZdNP during cuprizone feeding (W3 and W5), and 2 weeks after cuprizone removal (W5+2) using NestinCre^ERT2^:mTmG mice. Membrane GFP allows myelin segment visualization. Bottom row: high magnification images of GFP+ OLG (CC1+) are shown at each time point. Note that at W3, GFP+CC1+ cells do not yet exhibit the characteristic myelin segments as seen at W5+2. D: Phenotype of SVZdNP mobilized in the CC during cuprizone-induced demyelination. E: Relative densities of immature cells (neg cells) in medial vs lateral CC and in caudal vs rostral CC. Immature cells mobilized from SVZ are more numerous in lateral compared to medial CC as indicated by the mean ratio <1. Black bars show the means, n=4 mice. F-G: EGFR (F) and Nestin (G) immunolabeling in NestinCre^ERT2^:YFP mice exposed to cuprizone. Arrowheads point to YFP+ EGFR+ cells (G) and to YFP+ Nestin+ Olig2-cells. Cells in the white squares are shown at higher magnification. Scale bars A-C: upper panel: 200μm; lower panel: 20μm; F,G 20μm, zoom: 5 μm.

These results indicate that during cuprizone-induced demyelination, SVZdNP are recruited to the CC, quickly adopt an oligodendrocytic identity, but do not efficiently start to remyelinate axons until cessation of cuprizone exposure. Thus, early SVZdNP maturation in myelinating OLG is not sufficient to account for the differential myelin content along the CC at W3. We therefore hypothesize that rostral and lateral CC are protected from demyelination.

### A significant fraction of these cells remains undifferentiated in the demyelinated CC

Although SVZdNP quickly adopt an oligodendrocytic fate once recruited in the demyelinated CC, we nevertheless observed that a significant proportion of YFP^+^ SVZdNP remained negative for oligodendrocytic (CC1, Olig2), astrocytic (GFAP) or neuronal (DCX) markers (40.1 ± 9.9% and 27.4 ± 3,7% at W3 and W5 respectively, thereafter called “neg cells”) (Fig2D). The density of neg cells during demyelination is significantly higher in lateral compared to medial CC, with a ratio medial/lateral of 0.2±0.03 (p=0.02) (Fig2E). Some of these cells express EGFR or Nestin which are markers for C cells and neural progenitor cells (Fig2F,G). Thus, in a region that is protected from demyelination, we observe a significant proportion of SVZdNP that remain immature in the CC after cuprizone treatment.

### The density of activated microglial cells in the demyelinated CC is inversely proportional to SVZdNP mobilization

Since NSC have been shown to present immunomodulatory properties when grafted in injured brain (Kokaia et al. 2012), we hypothesized that SVZdNP that remain immature in the demyelinated CC could contribute to protect against demyelination through the modulation of the inflammatory micro-environment. We first investigated the presence of activated microglial cells. In control condition, microglial cells are homogeneously distributed along the CC (Suppl Fig1). After 3 weeks cuprizone exposure, microglial cells express CD68, a marker of activated microglia, (Fig3A) and their density is almost twice in medial compared to lateral CC (ratio medial/lateral = 1.81 ± 0.27; p=0,012) (Fig3A,B). This regionalized activation of microglia tends to persist at W5, although not significantly. Interestingly, areas enriched in SVZdNP show reduced numbers of activated microglial cells. Moreover, animals with the highest SVZdNP mobilization in the CC exhibited lowest microglial activation revealing a strong inverse correlation (R^2^= 0.978) between YFP+ cells density and activated microglia density (Fig3C). Thus, CC areas that are resistant to demyelination during cuprizone are rich in SVZdNP and poor in activated microglia.

**Figure 3:**
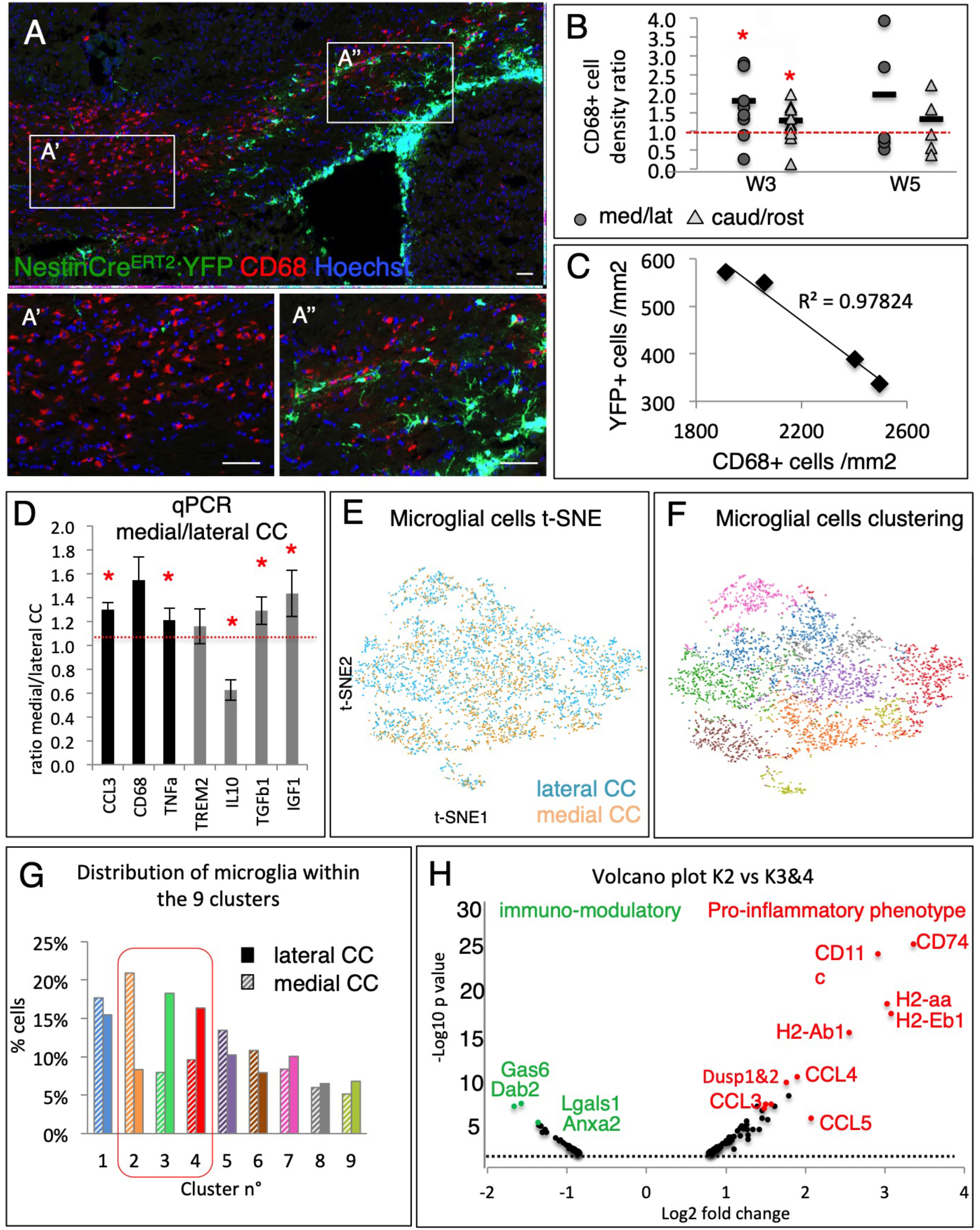
Inflammatory signature varies along the latero-medial axis in the demyelinated corpus callosum. A: Illustration of activated (CD68+) microglia in the CC of cuprizone-fed mice. White boxes in lateral (A’) and medial CC (A”) are shown at higher magnification. B: Quantification of CD68+ microglia density in medial relative to lateral CC (dark circles) and in caudal relative to rostral CC (light triangles). Black bars show the means, n=7 mice at W3 and 5 mice at W5. C: Inter-individual negative correlation between activated microglia and SVZdNP density in the CC of NestinCre^ERT2^:YFP mice fed for 3 weeks with cuprizone (n= 4 mice). D: RT-qPCR analysis of cytokine expression in medial compared to lateral CC. E: t-SNE representation after single cell RNA sequencing of microglial cells sorted from medial (2133 cells, in orange) and lateral CC (2304 cells, in blue) after 4 weeks cuprizone feeding. F: Unbiased automated clustering segregating microglial cells into 9 clusters, represented by different colors. G: distribution of microglial cells from medial and lateral CC in each of these 9 clusters. Cluster #2 is enriched in cells from medial CC, whereas clusters 3 & 4 are enriched in cells from lateral CC. H: Gene expression comparison between cluster 2 and clusters 3 & 4, represented as a volcano plot. In red, genes significantly upregulated in cluster 2 that identify M1 markers; in green, genes significantly upregulated in clusters 3 & 4 that identify M2 markers. Scale bar: 50 μm. See also Figure S1.

### Inflammatory signature varies along the latero-medial axis in the demyelinated CC

To test if SVZdNP could modulate microglial cell activation and thus exert protective functions we first examined if the different areas of the CC were associated with a particular inflammatory signature.

qPCR analyses in animals fed with cuprizone during 3 weeks showed that 2 pro-inflammatory cytokines, CCL3 and TNFa, but also of CD68, are significantly higher in the medial vs lateral CC (p=0.04 and 0.02 respectively; Fig3D). CD68 mRNA enrichment in medial CC confirmed the histological observations described above.

Among the anti-inflammatory cytokines tested, we found that IL10 is significantly lower in the medial CC (ratio medial vs lateral: 0,63 ±0.09, p=0.02; Fig3D). By contrast, IGF1 and TGFß1, two well-known immunomodulatory factors, are enriched in medial CC. These analyses suggest that during demyelination, CC areas with poor SVZdNP mobilization tend to exhibit an inflammatory profile more aggressive than those with high SVZdNP recruitment.

In order to compare the molecular signature of microglia in the medial vs lateral CC during demyelination, we FACS sorted microglia cells using CX3CR1GFP mice fed with cuprizone for 4 weeks and performed single-cell RNA-sequencing. We obtained the transcriptome of 2133 medial and 2304 lateral microglial cells.

The t-distributed stochastic neighbor embedding (tSNE) projection did not reveal a global segregation of the two samples (medial and lateral microglial cells) (Fig3E). However, the clustering, which ordered microglial cells in 9 clusters (Fig3F), pointed to differential distribution of microglia from medial and lateral CC for clusters 2, 3 and 4, the other clusters showing equivalent contribution (Fig3G). Cluster 2 was preferentially composed of medial microglia whereas clusters 3 and 4 were predominantly composed of lateral microglia. To further characterize these specific clusters (that represent 40.8% of all microglial cells analyzed), we compared the cluster enriched in medial microglia and the two clusters enriched in lateral microglia and examined the first 100 most significantly differentially regulated genes.

Almost all top genes enriched in cluster 2 (in which microglial cells from medial CC are predominant) are genes coding for pro-inflammatory cytokines (CCL3, CCL4, CCL5) or genes involved in antigen presentation (H2-aa, H2Ab1, H2-Eb1, CD74) characterizing the M1 phenotype (Fig3H). Conversely among the genes mostly enriched in clusters 3 and 4 (in which microglial cells from lateral CC are predominant) stand immunomodulatory genes characterizing the M2 phenotype (Gas6, Dab2, Lgals1, Anxa2)(Fig3H).

Thus, a large microglia population in the medial CC adopts a pro-inflammatory phenotype deleterious for myelin integrity and repair, while in lateral CC a microglia subpopulation exhibits an immunomodulatory phenotype associated to neuroprotection and regeneration.

### Identification of ligand-receptor couples involved in the dialog between SVZdNP and microglial cells

In order to identify SVZdNP proteins that may influence microglial phenotype during demyelination, we dissected the CC of NestinCre^ERT2^:RosaYFP mice fed during 4 weeks with cuprizone. YFP+ cells were FACS sorted, and analyzed by single-cell RNA-sequencing. t-SNE projection of 1931 SVZdNP transcriptomes followed by automated clustering ordered the cells in 6 distinct clusters (Fig4A). The enriched expression of typical markers allowed the identification of each clusters (Fig4B): 1-ependymal cells, 2-neuroblasts, 3-OPC, 4-proliferative neuroblasts, 5-neural progenitors, and 6-astrocytes. In order to identify ligand-receptor couples potentially implicated in the SVZdNP/microglia dialog, we focused on secreted extracellular molecules or membrane receptors known to exert immunomodulatory properties in Gene Ontology database. Among the 100 first upregulated genes we found Gas6 (Growth-arrest-specific protein 6) specifically enriched in cluster 1 identified as ependymal cells (16.7 fold, p=5.3E^−45^) and also present in the OPC cluster (2.1 fold, p=0.001) (suppl Fig2A). GAS6 is a soluble agonist of the TYRO3, AXL, MERTK (TAM) family of receptor tyrosine kinases identified to have anti-inflammatory and neuroprotective properties through the modulation of microglia activation (Gruber et al. 2014, Fourgeaud et al. 2016). AXL was indeed highly expressed in microglial cells in the demyelinated CC (suppl Fig2B). Consistent with the results of our approach, previous studies showed that GAS6-/- mice are more susceptible to cuprizone-induced demyelination specifically in the rostral part of the CC (Binder et al. 2008, Tsiperson et al. 2010, Ray et al. 2017). Our single-cell RNA-seq study thus suggests that ependymal cells and OPC could protect against demyelination by secreting GAS6.

**Figure 4:**
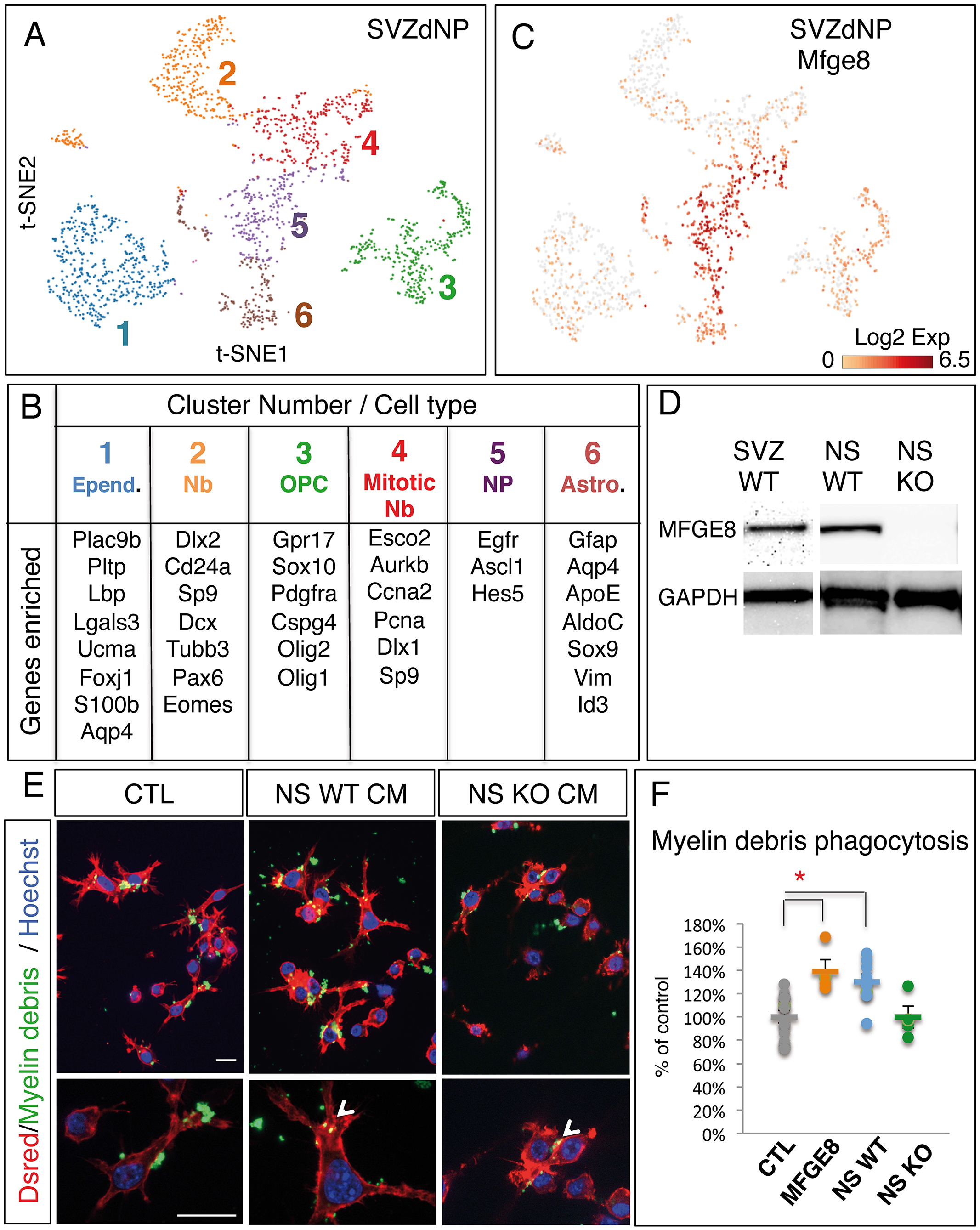
Molecular characterization of SVZdNP in demyelinated corpus callosum after 4 weeks cuprizone feeding and identification of MFGE8 as a candidate modulating microglial cells. A: t-SNE and automated clustering after single cell RNA sequencing of SVZdNP sorted from demyelinating CC. B: Identification of the 6 clusters generated by automated clustering using a list of genes enriched in each of these clusters. C: Expression levels of Mfge8 in SVZdNP showing enrichment in neural progenitors (cluster 5). D: Western blot confirming MFGE8 expression in SVZ and neurospheres derived from SVZ, but absence in neurospheres from MFGE8^−/-^ mice. E: Illustration of BV2 microglial cells stably expressing DsRed, cultured in presence of CFSE-labeled myelin debris, and treated or not with neurosphere (NS)-conditioned medium (CM). Scale bars: 20 μm. F: Quantification of myelin debris phagocytosis by microglial cells in vitro after addition of rmMFGE8 or of neurosphere-conditioned medium from wild-type (WT) or MFGE8^−/-^ mice (KO).

Immature neural progenitors (cluster 5 in Fig4A,B) are the most likely population among SVZdNP to be endowed with immunomodulatory properties. The 3 most significantly enriched genes in cluster 5 (Fig4B,C) are Egfr, (2.7 fold; p= 1.41E^−16^) Mfge8 (2.7 fold; p= 5.87E^−13^) and Ascl1 (2.1 fold; p= 2.80E^−11^). EGFR (epidermal growth factor receptor) and ASCL1 (achaete-scute homolog1) are indeed commonly used markers for C cells. MFGE8 (milk fat globule-epidermal growth factor-8) and its receptor β3 integrin (Itgb3) are known to induce phagocytosis of apoptotic cells and to act as “endogenous protective factors” in response to various brain lesions (Deroide et al. 2013, Liu et al. 2015, Xiao et al. 2018). We found β3 Integrin expressed in most of the microglial cells present in the demyelinated CC (Suppl Fig 2C).

Because of MFGE8 / integrin ß3 mode of action, we hypothesized that MFGE8 secreted by SVZdNP may promote myelin debris endocytosis by microglial cells which is crucial for inflammation resolution. Indeed, the proportion of BV2 microglial cells engulfing CFSE-labeled myelin debris in vitro increased by 39% (p=0.03) after addition of MFGE8 in the culture medium (Fig4E).

Then, to demonstrate that SVZ progenitors secrete MFGE8 and promote myelin debris phagocytosis by microglial cells, we cultured SVZ cells as neurospheres from WT and MFGE8 ^−/-^ mice and collected the conditioned medium. SVZ cells from WT mice cultured as neurospheres secrete MFGE8 (Fig4D), and their conditioned medium significantly enhances myelin debris phagocytosis (p=0.01; Fig4E,F). By contrast, conditioned medium from neurospheres derived from mutant mice lacks MFGE8 (Fig4D) and fails to promote myelin debris phagocytosis (Fig4E,F).

Altogether these results therefore support the hypothesis that SVZdNP, mobilized upon cuprizone treatment, protect against CC demyelination by reducing inflammation and promoting myelin debris phagocytosis via GAS6 and MFGE8 secretion.

## DISCUSSION

Neural stem cells in SVZ produce neuroblasts and OPC that contribute to myelin repair after lesion (Menn et al. 2006; Jablonska et al. 2010; El waly et al. 2018; Xing et al. 2014, Brousse et al. 2015). A few studies suggested that SVZdNP respond to demyelination and migrate to the lesion site, but fail to produce myelin (Kazanis et al. 2017; Serwanski et al. 2018). It was also proposed that SVZdNP are dispensable for myelin repair but protect neurons from degeneration (Butti et al. 2019). In the present study, we show that in addition to OLG replacement, SVZdNP contribute to minimize demyelination by modulating microglial activity and promoting myelin debris phagocytosis.

In accordance with previous studies (Steelman et al. 2012, Chrzanowski et al. 2019, Zhang et al. 2019), we observe a regionalized sensitivity to cuprizone-induced demyelination, with reduced myelin loss in rostral and lateral CC. This could either indicate protection from demyelination or early remyelination. Our experiments do not support the “remyelination hypothesis”: indeed, myelin segments produced by SVZdNP (visible using mTmG reporter) are rare in CC at early time points. Beside, higher myelin content in rostral and lateral areas cannot be attributed to pOPC since they preferentially contribute to remyelination in caudal and medial CC (Xing et al. 2014, Brousse et al. 2015).

Numerous studies revealed that grafted NSC exhibit a “homing” behavior and contribute to regeneration via bystander effects independent of cell replacement (for review see (Kokaia et al. 2012). They create niches of neural progenitors that trigger apoptosis of infiltrating T cells (Pluchino et al. 2005) and modulate the expression of inflammatory transcripts in microglia (Cusimano et al. 2012). Whereas the immunomodulatory and trophic properties of grafted stem cells are now well documented (for review see Ottoboni et al., 2015), such possible functions of endogenous SVZdNP have never been investigated in injured brain. Our work provides the first evidence for immunomodulatory functions of endogenous SVZdNP in a demyelination context. We show that CC areas enriched in SVZdNP display reduced densities of activated microglial cells, and host subpopulations of microglial cells characterized by an immunomodulatory profile. We do not exclude that SVZdNP may act both locally after mobilization to the lesioned site, and remotely, from their niche. In our RNA-seq experiment, the presence of numerous ependymal cells and neuroblasts suggests that we probably also collected SVZdNP from the niche in addition to SVZdNP mobilized in the CC, due to the apposition of the dorsal wall of the SVZ to the CC.

Microglia are key players in the demyelination/remyelination process, with both deleterious and beneficial properties. Recent studies using single-cell RNA-sequencing revealed multiple microglial phenotypes, complicating the simplistic dichotomous view of M1 versus M2 phenotype (Keren-Shaul et al. 2017). Efficient remyelination requires the death of pro-inflammatory microglia and repopulation by pro-regenerative microglia (Miron et al. 2013; Lloyd et al., 2019). A prerequisite for inflammation resolution and remyelination is myelin debris phagocytosis by microglia (Kotter et al. 2006, Lampron et al. 2015). In a model of inflammatory demyelinating disease, the experimental autoimmune encephalomyelitis, microglial phagocytic activity is correlated to functional recovery (Yamasaki et al. 2014). In this study, we propose that at least two ligand/receptor couples, GAS6/AXL and MFGE8/ß3 integrin, could be involved in immunomodulatory effects of endogenous SVZdNP via microglia modulation. We found GAS6 enriched in ependymal cells and SVZ-derived OPC, and AXL widely expressed by microglial cells. Interestingly, Gas6-/- mice exposed to cuprizone exhibit increased demyelination specifically within the rostral CC compared to wild type mice (Binder et al. 2008, Tsiperson et al. 2010, Ray et al. 2017). Our results indicate that the SVZ niche is a source of GAS6 and thus highlight the protective function of endogenous SVZdNP.

MFGE8 was found highly enriched in neural progenitors and ß3 integrin expressed at low level in most microglial cells. MFGE8 has been shown to recognize phosphatidylserine “eat me” signal and binds to ß3 integrin expressed by macrophages mediating phagocytic clearance of apoptotic cells (Fadok et al. 2000, Hanayama et al. 2002). Recent studies suggest that MFGE8 could act as a protective factor endowed with immunomodulatory properties (Tan et al. 2015; Xiao et al. 2018) by inhibiting M1 microglia polarization (Shi et al. 2017; Li et al. 2019). Here we show that MFGE8 or SVZdNP-conditioned medium enhance myelin debris phagocytosis by microglial cells in vitro, the effect of conditioned medium being lost when SVZdNP are derived from MFGE8-/- mice.

These results suggest that endogenous SVZdNP activated by demyelination insult contribute to reduce inflammation and promote myelin debris phagocytosis via GAS6 and MGFE8 secretion, thus protecting against extensive demyelination and creating a local environment more favorable to the regenerative process.

## Acknowledgments

We thank C. Maurange, C. Faivre Sarrailh and L. Kerkerian Le Goff for critical reading of the manuscript. This work was funded by CNRS, Aix-Marseille University, the Fondation pour la Recherche Médicale (DEC 20140329501) and the ARSEP fondation. We also acknowledge France-bioimaging/PICSL infrastructure (ANR-10-INSB-04-01) and NeuroSchool/nEURo*AMU (ANR-17-EURE-0029). BB was funded by ANR and an ARSEP fellowship.

## Author Contributions

BB performed most experiments and analyzed the data, with the help of KM; FD performed bioinformatics analyses of single cell RNA sequencing; PD and MC directed the study and wrote the manuscript.

## METHODS

All experimental and surgical protocols were approved by the ethic committee for animal experimentation (reference 2016071112151400).

### Animals

NestinCre^ERT2^ (Lagace et al. 2007) transgenic mice were used to trace SVZdNP. Heterozygous Cre mice were crossed with homozygous or heterozygous R26R-YFP (Srinivas et al. 2001) or mTmG (Muzumdar et al. 2007) reporters to generate double-heterozygous offspring for cell lineage analysis. CX3CR1GFP (Jung et al. 2000) heterozygous transgenic mice were used to label and sort microglial cells. PLPGFP (Le Bras et al. 2005) transgenic mice were used to image and quantify myelin in the CC. C57BL/6 (Janvier lab) were used for western blot experiment.

### Tamoxifen injection and demyelination

Six-week-old NestinCre^ERT2^:R26R-YFP or NestinCre^ERT2^:mTmG mice were injected for 5 consecutive days with tamoxifen (180 mg/kg) to induce recombination and cell labeling. To induce demyelination, cuprizone (Sigma) treatment (0.2% in food) started 2 weeks after the end of tamoxifen injections and lasted for 5 weeks. Mice were sacrificed after 3 (W3), 4 (W4), 5 (W5) weeks of cuprizone treatment or 2 (W5+2) and 4 (W5+4) weeks after cuprizone removal.

### Tissue preparation and immunohistochemistry

After intracardiac perfusion with 4% paraformaldehyde, brains were removed, postfixed for 2 h and cryoprotected in sucrose 20% overnight. After OCT inclusion brain were cut on cryostat on serial coronal sections 20 μm thick. We used the following primary antibodies: chicken anti-GFP (1/500; Aves Labs), rabbit anti-Olig2 (1/500; Chemicon), mouse anti-GFAP (1/1000; Sigma), goat anti-DCX (1/250; Santa Cruz), anti-Nestin (1/100; Millipore), anti-EGFR (1/50; Cell Signaling), rat anti-CD68 (1/200 AbCam).

### Microscopy and quantification

Images were captured with a Zeiss Apotome system (20× objective). CC was analyzed in two rostro-caudal locations: above the rostral SVZ (Bregma +0.5 to +1) and caudal to the fornix (Bregma −0.3 to −0.8) (Fig. 1C) and two medial-lateral location (midline and above the lateral ventricles).

Sections from at least three mice were analyzed at each time point. For each mouse, three sections in each location were analyzed. The area of interest was delineated and the cells of interest were counted using Zen software (Zeiss). For quantification of GFP signal, exposure time was fixed for all images and analysis was done using ImageJ software. Within each mouse, GFP signal was compared in caudal vs rostral areas and in medial vs lateral areas of the CC, and was thus expressed in ratios.

### Generation of CC single-cell suspension

For single-cell RNA-seq experiments or for RTqPCR, SVZdNP and microglial cells were isolated from demyelinated CC from NestinCre^ERT2^:R26R-YFP or CX3CR1GFP mice respectively. Four to five mice were deeply anesthetized and transcardially perfused with 25 ml ice cold phosphate buffer saline (PBS) (Sigma). Brains were immediately extracted and collected in ice cold Hanks Balanced Salt Solution (HBSS) without Ca2+ and Mg2+ (Invitrogen). Brains were cut into 500μm slices with a vibratome (Microm). For SVZdNP single-cell RNA-seq, CC was microdissected and cut into 3-4 pieces to facilitate dissociation. For microglia single-cell RNA-seq medial and lateral CC from each mouse were separately microdissected to obtain two distinct samples. Cortex from P1 to P7 wild type neonates were used as ballast to avoid losing cells during successive centrifugations. Enzymatic and mechanical dissociations were done using the Neural Tissue Dissociation Kit (P) (Miltenyi) and the automated gentle MACS (37C-NTDK-1 program) following manufacturer’s instructions. To proceed for myelin debris removal we used Miltenyi magnetic beads. Finally, cells were resuspended in 500 μl of phosphate saline buffer (PBS) with 0.5% BSA.

### FACS

YFP+ or GFP+ cells, respectively in NestinCre^ERT2^:R26R-YFP and in CX3CR1GFP mice, were FACS sorted (BD FACS ARIA, CIPHE) and collected into a coated tube with sorting buffer (PBS, 0.5%BSA) for subsequent single-cell RNA-seq experiment or collected into a tube containing lysis buffer for RNA extraction. For single-cell RNA-seq experiments, 5000 cells/sample were collected in a maximum volume of 20μl.

### Single-cell library generation and sequencing

Single-cell library generation (10X Chromium) and sequencing (NextSeq 500) were performed by the HalioDX Company (Luminy, Marseille). Clustering and gene expression analysis was done using Cell Ranger and Loupe Cell Browser software developed by 10X Genomics.

### qPCR experiment

Total RNA was extracted from tissues (CC) or cells (microglial GFP+ cells sorted by FACS) using the RNeasy Mini Kit (Qiagen). After reverse transcription using Superscript First-Strand Synthesis System (Invitrogen), cDNA, primers, and the Syber Green Master Mix (Life Technologies) were mixed as instructed by the manufacturer, and RT-qPCR and melting curve analysis were performed on CFX96 (BioRad). Primers used for a variety of pro- and anti-inflammatory genes are the following: Ccl3 forward TGC CCT TGC TGT TCT TCT CT Ccl3 reverse GTG GAA TCT TCC GGC TGT AG; Cd68 forward GAC CTA CAT CAG AGC CCG AGT Cd68 reverse CGC CAT GAA TGT CCA CTG; Tnfa forward TGC CTA TGT CTC AGC CTC TTC RTnfa reverse GAG GCC ATT TGG GAA CTT CT; Trem2 forward ACC CAC CTC CAT TCT TCT CC Trem2 reverse GAT GCT GGC TGC AAG AAA CT; Il10 forward ACC TGC AGT GTG TAT TGA GTC TG Il10 reverse CCC TGG ATC AGA TTT AGA GAG C; Tgfß1 forward CAA TTC CTG GCG TTA CCT TG Tgfß1 reverse AGA CAG CCA CTC AGG CGT AT; Igf1 forward CTC ACC TTC ACC AGC TCC Igf1 reverse CAC GAA CTG AAG AGC ATC CA; Gapdh forward GGC CTT CCG TGT TCC TAC Gapdh reverse TGT CAT CAT ATC TGG CAG GTT

### Western blot

After brain extraction, 400μm slices were performed with vibratome (Microm) and SVZ, was dissected in cold HBSS (Invitrogen). Tissues were instantly frozen on dry ice. Tissues (SVZ or neurospheres) were crushed and sonicated in lysis buffer (1 ml /100 mg tissue; 50 mM Tris-HCl pH 7.5, 150 mM NaCl, 1 mM EDTA, 1 mM PMSF, anti-proteases, 1% Triton) and supernatant collected after centrifugation (1 h, 14 000 rpm, 4°C). Supernatant proteins (20 to 40μg load, quantified by Pierce BCA protein assay kit) were resolved by SDS/PAGE on 4-12% gels (Invitrogen) and blotted onto PVDF membranes. For Western blotting, membranes were blocked for 1 h in TBS-T (0.02 M Tris pH 8, 0.5 M NaCl, 0.1% Tween-20) containing 4% dehydrated half-creamed milk, and incubated (overnight, 4°C) rabbit anti-MFGE8 (1/1000, Euromedex), then with anti-mouse horseradish peroxidase-conjugated antibodies (1/3000; 2 h; Jackson ImmunoResearch) and revealed with ECL substrate (Lumi-Light Western Blotting Substrate; Pierce chemical company). Chemiluminesce was registered under ECL Imager Chemi-Smart system (Fisher Bioblock).

### Primary culture of SVZ grown as neurospheres

Neurospheres were produced as previously described in Durbec and Rougon (2001) (Durbec and Rougon 2001) from P2-P5 newborn mice, in defined media supplemented with 2% B27 and 20ng/ml EGF and bFGF (R&D System). Neurospheres were passaged once and conditioned medium was recovered before the second passage, filtered through 0.2 *μ*m filters (Acrodisc serynge filters; Cell Science) and stored at −80°C before use.

### Myelin debris phagocytosis by microglia cells

The preparation of fluorescent-labeled myelin debris was performed as described in (Rolfe et al. 2017). Briefly, brains from 10 adult mice were slices into small pieces, homogenized in sucrose 0.32M and then centrifuged through sucrose 0.83M (100,000xg 45 min at 4°C). Myelin debris were collected at the interface between the 2 sucrose concentrations. After 2 successive Tris buffer rinses/centrifugation steps, myelin debris pellets are finally resuspended in PBS (100mg/ml) and stored at −80°C. Before use, myelin debris are fluorescently labeled by incubation in carboxyfluorescein succinimidyl ester (CFSE) 50 μM for 30 min at room temperature, washed (PBS glycine 100mM) 3 times, and after final centrifugation (14800xg 10 min 4°C), fluorescent myelin debris were resuspended in sterile PBS (100 mg/ml).

BV2 cells (mouse brain microglial cells) were seeded at 200,000 cells/ml in 4-wells Lab-Tek (Dutscher). 24 h later, 5 μl mMFGE8 (500 ng/ml; Santa Cruz) was added to the culture medium (or PBS for control), and 1 hr after MFGE8 addition, CFSE-labeled myelin debris (1 mg/ml) were added in the culture. Cells were fixed 3h later with 4% paraformaldehyde for 30 min, rinsed, and nuclei were labeled by 5 min incubation in Hoechst. 10 pictures were taken by well at x20 magnification, and the proportion of phagocyting microglia was quantified.

The same experiment was performed using neurosphere-conditioned medium instead of MFGE8 treatment (and not conditioned neurosphere defined medium as control).

To obtain neurosphere-conditioned medium, neurospheres were cultured as previously described (Durbec and Rougon, 2001). Briefly, SVZ were dissected from wild type or MFGE8 KO neonate mice, dissociated (trypsine 5 mg/ml) and maintained in defined medium supplemented with 2% B27 and 20 ng/ml EGF and bFGF. Conditioned medium was collected 4 days after the third passage, filtered and stored at −80°C.

### Statistical analyses

Non-parametric Mann & Whitney and Wilcoxon tests were used for 2-group comparisons (unpaired and paired samples respectively). For multiple group comparison, ANOVA by score permutation was performed followed by Kruskal Wallis, using the StatXact software (Cytel Studio). Differences were considered as significant for p<0.05 (*) and highly significant for p<0.01 (**).

## SUPPLEMENTAL INFORMATION TITLES AND LEGENDS

**Supplementary Figure 1: In healthy mice not exposed to cuprizone diet, microglial calls are evenly distributed along the CC**.

A: Illustration of microglial labeling in healthy CX3CR1-GFP mice. White boxes in medial (A’) and lateral CC (A”) are shown at higher magnification. B: Quantification of GFP+ cells in the CC of CX3CR1-GFP healthy mice.

**Supplementary Figure 2: Identification of GAS6/AXL as a ligand/receptor couple potentially involved in the dialog between SVZdNP and microglial cells.**

A: Expression levels of Gas6 in SVZdNP showing enrichment in ependymal cells and to a lesser extent OPC. B: Expression levels of Axl, a Gas6 receptor, in microglial cells during cuprizone-induced demyelination. C: Expression levels of Itb3, a receptor for MFGE8, in microglial cells during cuprizone-induced demyelination

