## supplementary figures for "Endogenous neural stem cells modulate microglia and protect from demyelination"

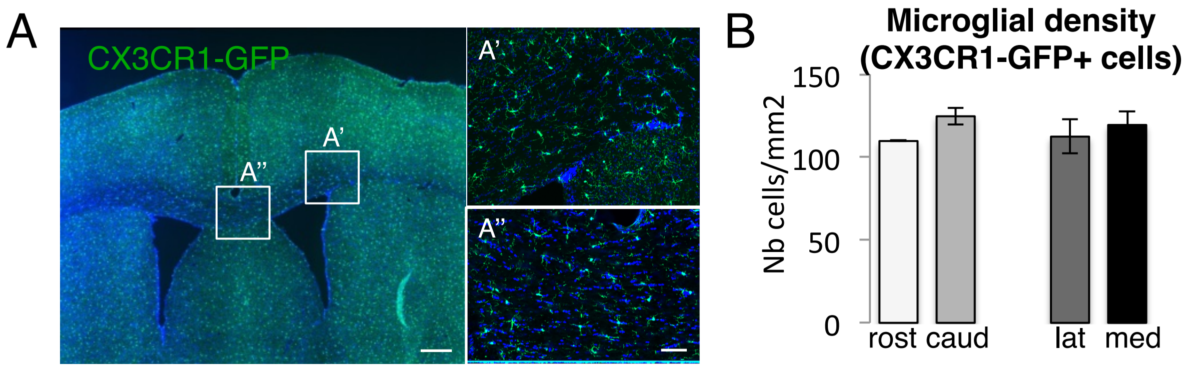


**Supplementary Figure 1: In healthy mice not exposed to cuprizone diet, microglial calls are evenly distributed along the CC**.

A: Illustration of microglial labeling in healthy CX3CR1-GFP mice. White boxes in medial (A') and lateral CC (A") are shown at higher magnification. B: Quantification of GFP+ cells in the CC of CX3CR1-GFP healthy mice.

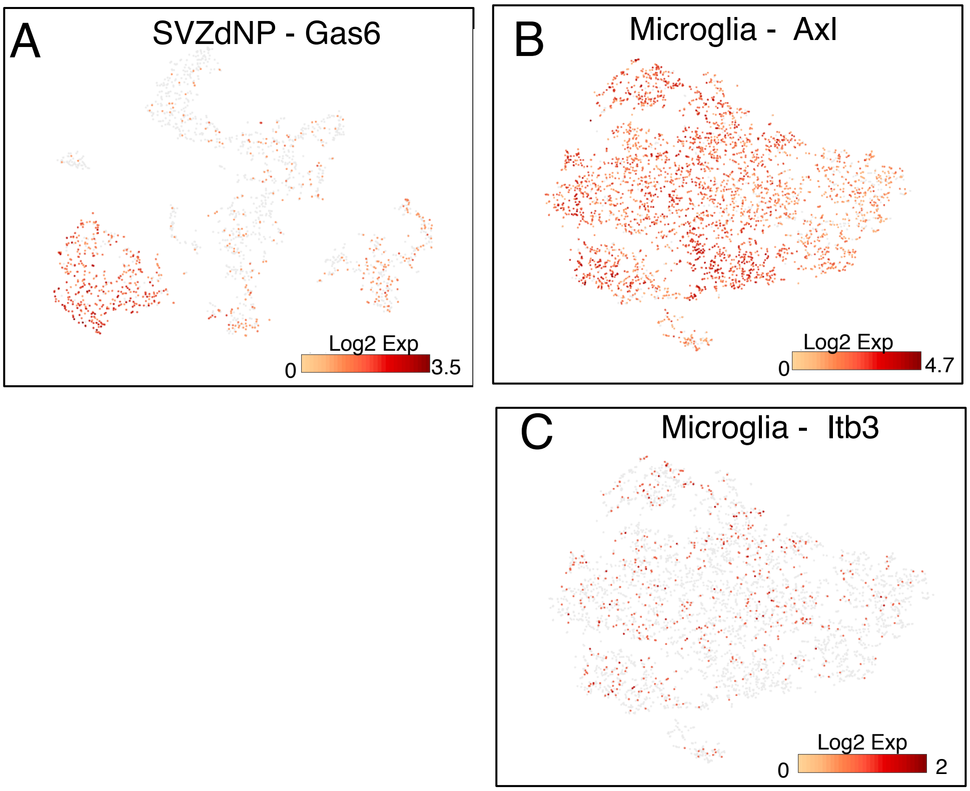


**Supplementary Figure 2: Identification of GAS6/AXL as a ligand/receptor couple potentially involved in the dialog between SVZdNP and microglial cells.**
A: Expression levels of Gas6 in SVZdNP showing enrichment in ependymal cells and to a lesser extent OPC. B: Expression levels of Axl, a Gas6 receptor, in microglial cells during cuprizone-induced demyelination. C: Expression levels of Itb3, a receptor for MFGE8, in microglial cells during cuprizone-induced demyelination
